# Mechanical Signaling Regulates DNA Methylation to Maintain Muscle Stem Cell Quiescence

**DOI:** 10.64898/2026.05.15.725521

**Authors:** Pukana Jayaraman, Paige R. Deltener, Justice Paintsil, Christian O. Abosede, Atreyi Ghatak, Sarah Abrahamson, Chris W.D. Jurgens, Damien Parrello, Susan Eliazer

## Abstract

Skeletal muscle stem cells (MuSCs) reside within a mechanically dynamic niche where they integrate biophysical and biochemical cues to maintain quiescence. Here, we show that substrate stiffness and RhoA-dependent signaling regulate MuSC fate. MuSCs cultured on soft matrices or depleted of RhoA exhibit altered morphology, diminished actomyosin organization, and undergo premature activation. Loss of *RhoA* reshapes the DNA methylation landscape, leading to widespread changes in gene expression and alternative splicing. Dnmt3a was among the genes transcriptionally downregulated following loss of RhoA signaling. Mechanistically, RhoA maintains *Dnmt3a* expression by promoting SP1 occupancy at its promoter. Importantly, loss of *Dnmt3a* in quiescent MuSCs is sufficient to drive activation, identifying *Dnmt3a* as a key epigenetic effector downstream of mechanical signaling. Together, these findings define a mechanotransduction-epigenetic axis in which RhoA maintains stem cell quiescence by preserving DNA methylation programs through Dnmt3A.

## INTRODUCTION

Adult stem cells rely on signals from their microenvironment to keep them in a quiescent state.^1^ In the skeletal muscle, stem cells (MuSCs) occupy a defined niche between the plasma membrane of the muscle fiber and the basal lamina.^2,3^ The adult MuSCs remain in a quiescent, non-dividing state, until injury triggers their activation.^4,5^ The activated MuSCs then proliferate, differentiate, and repair the injured muscle tissue.^6,7^ The maintenance of this quiescent state is essential for the long-term preservation of the MuSC population. Loss of quiescence leads to stem cell depletion and impaired regeneration.^4^

MuSCs are continuously exposed to the mechanical features of the microenvironment, including the contractility of the muscle fibers, the stiffness of the extracellular matrix, and compressive forces imposed by the surrounding tissue architecture.^8,9^ These mechanical inputs are sensed through integrins, cadherins, and mechanosensitive ion channels that influence stem cell fate and behavior.^10–12^ MuSCs are mechanosensitive and can sense and respond to mechanical cues. Mechanical confinement or compression of MuSCs *in vitro* alters the stem cell state.^13,14^ We have previously shown that niche factor Wnt4, a molecule secreted by the muscle fibers, maintains MuSC quiescence by regulating its mechanical properties.^15^ Wnt4 functions non-cell autonomously through the RhoA-ROCK signaling pathway to inhibit Yes Associated Protein 1 (YAP1), a well-known mechanotransducer and an activation marker, in MuSCs.^15–17^ How these mechanical inputs are converted into cell-intrinsic programs that maintain MuSC quiescence remain unknown.

A candidate mechanism that enforces stable cellular states is DNA methylation, an epigenetic mechanism that preserves cell identity.^18,19^ DNA methylation regulates gene expression without altering the underlying DNA sequence. It occurs through the addition of a methyl group to the 5’ position of cytosine residues (5mC) within CpG dinucleotides.^20,21^ Hypermethylated string of CpG nucleotides (islands) in promoter regions may cause gene silencing, whereas hypermethylation in gene bodies cause increased gene expression.^22–24^ This epigenetic mark is maintained by DNA methyltransferase, Dnmt1 and established *de novo* by Dnmt3A and Dnmt3B. The removal of these marks is mediated by Tet family enzymes, Tet1, Tet2, and Tet3.^25^ The functional importance of DNA methylation is well established in stem cells. For instance, in hematopoietic stem cells (HSCs) it balances long-term self-renewal with lineage commitment.^26,27^ Similarly, in MuSCs, DNA methylation regulates their proliferation and differentiation.^28–32^

Beyond its canonical role in transcriptional regulation, DNA methylation has also been implicated in regulating alternative splicing of RNA.^33,34^ This is particularly relevant to MuSCs as recent work suggests that alternative splicing contributes to the maintenance of quiescence.^35,36^ Despite the importance of DNA methylation in maintaining cell identity, its role in quiescent MuSCs remains largely unexplored. Furthermore, the mechanisms by which mechanical signaling maintains DNA methylation states in the genome of quiescent stem cells remains unknown.

Here, we show that RhoA-mediated mechanical signaling maintains MuSC quiescence by establishing a DNA methylation landscape that governs both transcriptional and post-transcriptional programs. Mechanistically, RhoA activity promotes transcription factor SP1 occupancy at the *Dnmt3a* promoter, sustaining *Dnmt3a* expression required for MuSC quiescence. Together, these findings connect niche-derived mechanical cues to epigenetic control of stem cell state, preventing premature activation.

## RESULTS

### Changes in cellular mechanoproperties drive stem cell fate decisions

To test whether mechanical inputs directly regulate MuSC fate, we altered extracellular matrix stiffness and assessed early molecular and morphological changes. MuSCs that are CD31^-^, CD45^-^, Sca1^-^, Vcam1^+^, and α7-integrin^+^ were sorted from wild-type (WT) mice. This surface marker profile excluded endothelial cells (CD31-), hematopoietic cells (CD45-), and fibro-adipogenic progenitors (Sca1-), while enriching for MuSCs expressing Vcam1 and α7-integrin. These freshly isolated MuSCs were plated immediately onto matrices of defined stiffness (0.5, 16, or 32 kPa), with 16 kPa approximating physiological stiffness,^37^ and analyzed after 10 hours (Figure 1A). Cell size increased progressively with stiffness, as observed by increased cell area (Figure 1B, 1C) and reduced circularity (Figure 1D). In contrast, nuclear size did not scale with cell area but instead, MuSCs cultured at physiological stiffness displayed a significantly larger nuclear area than cells on either softer or stiffer matrices (Figure 1E), suggesting that nuclear morphology is optimized at physiological stiffness.

**Figure 1.**
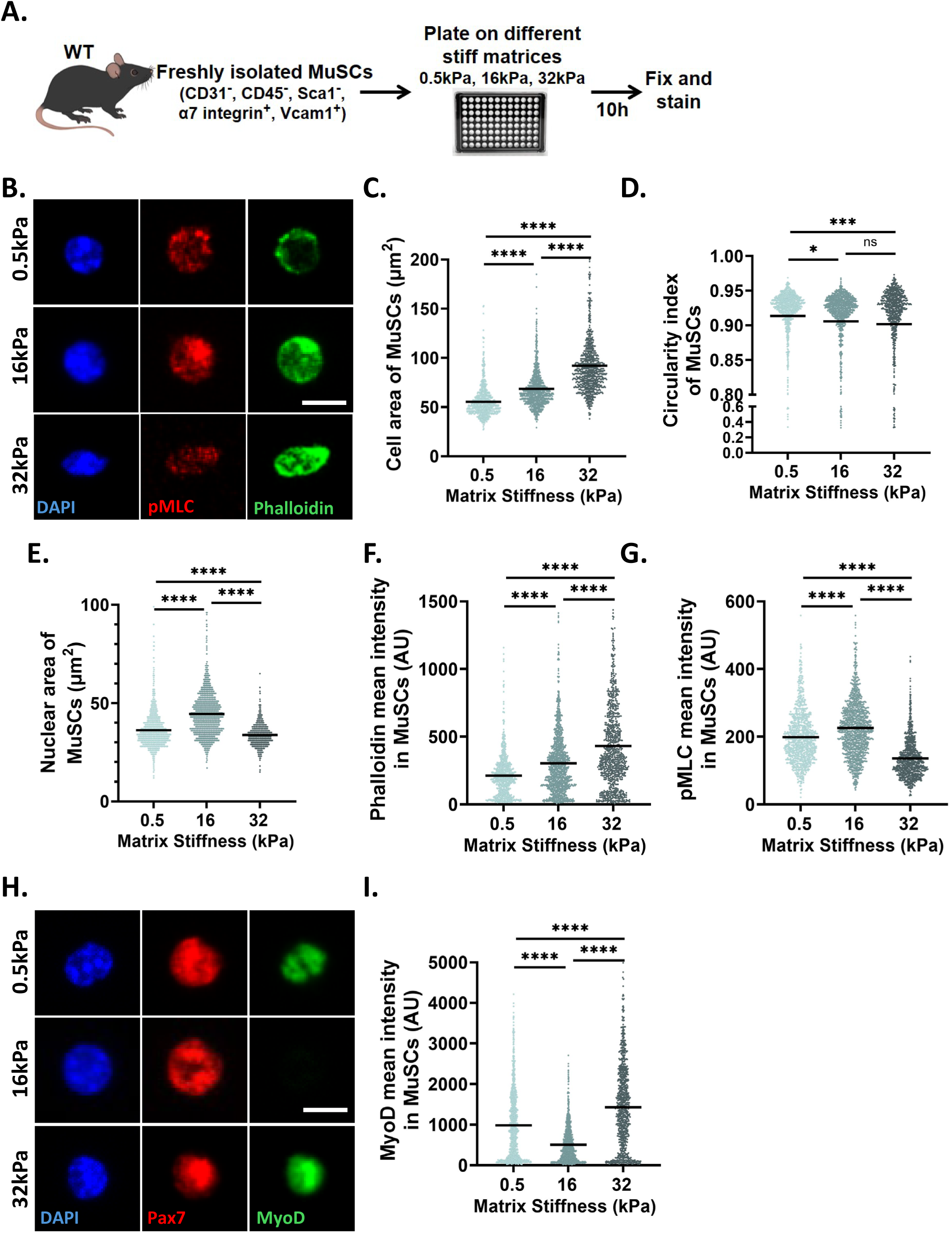
Substrate stiffness regulates MuSC mechanics and fate. (A) Schematic of the experimental design. (B) Representative images of FACS isolated WT MuSCs cultured on matrices of different stiffness, stained for pMLC and Phalloidin. (C - G) Quantification of cell area (C), cell circularity (D), nuclear area (E), Phalloidin (F), and pMLC intensity (G) in isolated MuSCs plated on substrates of varying stiffness for 10 hours (n = 4). (H and I) Representative images (H), and quantification of MyoD intensity (I) in MuSCs plated on different stiffness matrices for 10 hours (n = 4). Each dot represents an individual MuSC; the line indicates the median. Statistical significance was determined using an unpaired two-tailed Student’s t test.*p<0.05, ***p < 0.001, ****p < 0.0001, ns- not significant; scale bars, 5µm in (B) and (H).

F-actin levels, quantified by Phalloidin staining, increased with substrate stiffness, paralleling changes in cell area (Figure 1B, 1F). In contrast, actomyosin contractility, measured by the phosphorylation of myosin light chain (pMLC) at serine 19,^38^ peaked at physiological stiffness and was reduced under both softer and stiffer conditions (Figure 1B, 1G). This response followed the same pattern as the changes in nuclear size, suggesting that physiological stiffness sets an optimal condition that maintains quiescence with balanced cellular tension.

We next asked whether these mechanically defined states correlate with stem cell fate. MuSC activation, assessed by MyoD expression,^39^ was elevated on soft and stiff substrates relative to physiological stiffness (Figure 1H, 1I). These findings indicate that substrates recapitulating the physiological stiffness of native muscle promote maintenance of MuSC quiescence, whereas shifts toward either softer or stiffer mechanical environments bias MuSCs toward activation.

### RhoA-dependent cytoskeletal tension maintains the quiescent MuSCs *in vivo*

To test whether the mechanical state of the MuSCs at physiological stiffness is intrinsically controlled, we reduced RhoA signaling, a small G protein that functions as a central regulator of actin organization and actomyosin contractility.^40,41^ Since the complete loss of RhoA induces MuSCs to differentiate into Myogenin^+^ cells and fuse into the adjacent muscle fibers,^42^ we used an inducible genetic strategy in which a single RhoA allele (Rho^fl/+^) was deleted in MuSCs using tamoxifen inducible Pax7^CreER^.^43,44^ In this RhoA knockdown model, there is an increased number of Pax7^+^ MuSCs at homeostasis and they are not lost to differentiation.^15,42^ This mouse line is hereafter referred to as MuSC-RhoA^fl/+^, with Cre-negative littermates serving as controls (Figure 2A). Tamoxifen treatment of control and MuSC-RhoA^fl/+^ mice for seven consecutive days followed by a 30-day recovery period resulted in a 50% reduction of RhoA transcripts (Figure 2B) and a decrease in active RhoA protein levels (Figure 2C, 2E), confirming effective downregulation of the pathway.

**Figure 2.**
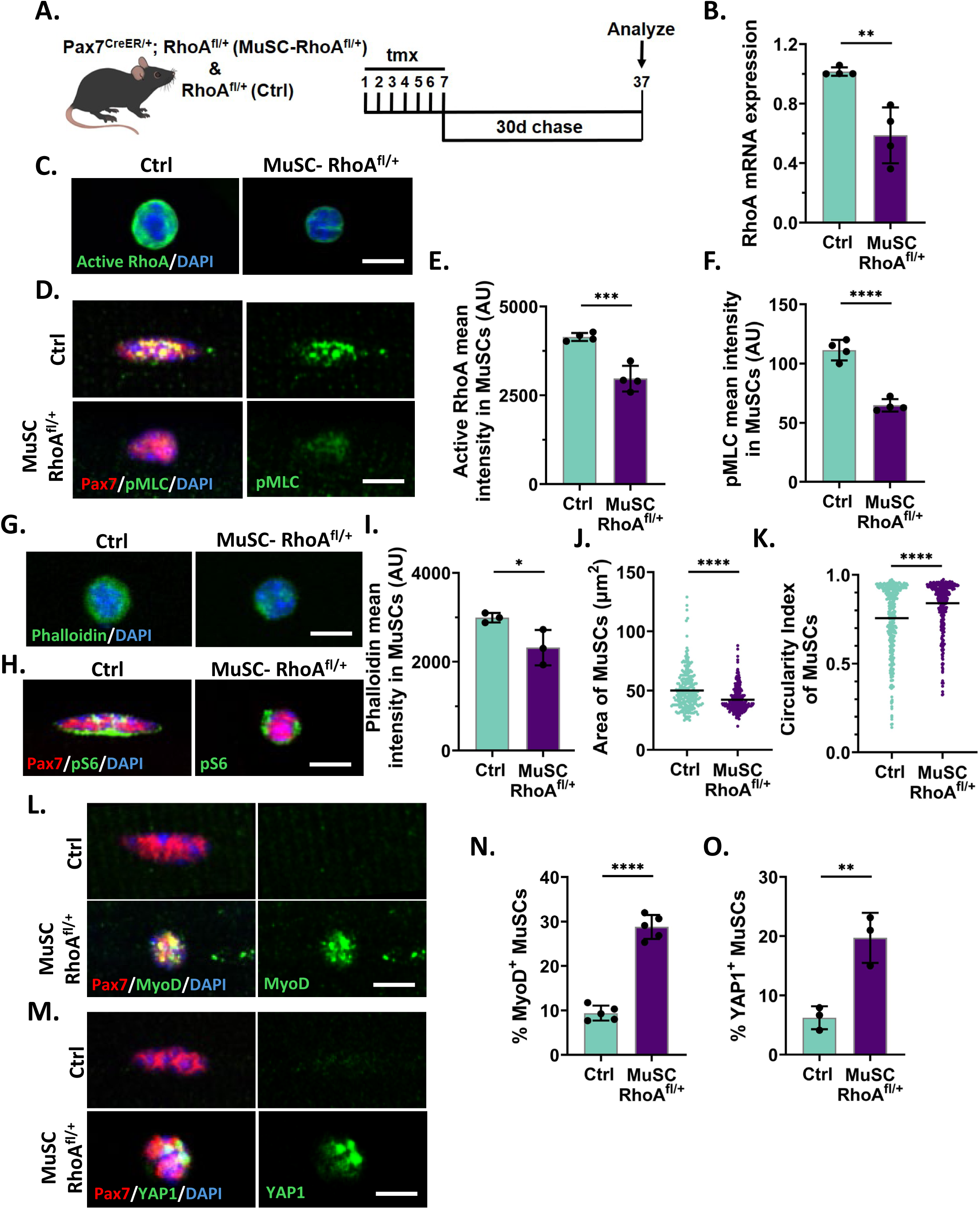
Mechanical signaling through RhoA maintains MuSC quiescence. (A) Schematic representation of the experimental design. (B) RhoA mRNA expression in freshly isolated MuSCs from control and MuSC-RhoA^fl/+^ mice 30 days after tamoxifen injection. Expression levels were normalized to GAPDH, and fold change relative to control samples was determined using the ΔΔCt method (n = 4). (C and E) Representative images (C) and quantification (E) of active RhoA intensity in FACS isolated MuSCs from control and MuSC-RhoA^fl/+^ mice (n = 4). (D and F) Representative images (D) and quantification (F) of pMLC levels in MuSCs on isolated myofibers from control and MuSC-RhoA^fl/+^ mice (n = 4). (G and I) Representative images (G) and quantification (I) of Phalloidin intensity in FACS isolated MuSCs from control and MuSC-RhoA^fl/+^ mice (n = 3). (H, J, and K) Representative images of MuSCs stained for pS6 (H) with quantification of cell area (J) and circularity (K) in MuSCs on control and MuSC-RhoA^fl/+^ myofibers (n = 3). (L and N) Representative images (L) and quantification (N) of MyoD intensity in MuSCs on control and MuSC-RhoA^fl/+^ myofibers (n = 5). (M and O) Representative images (M) and quantification (O) of Yap1 intensity in MuSCs on control and MuSC-RhoA^fl/+^ myofibers (n = 3). Each dot in bar graphs represents an individual mouse; each dot in dot plots represents an individual MuSC, with the central line indicating the median. Statistical significance was determined using an unpaired two-tailed Student’s t test. Error bars, mean ± SD; *p<0.05, **p < 0.01, ***p < 0.001, ****p < 0.0001; scale bars, 5µm in (C, D, G, H, L, and M).

We next examined the mechanical consequences of reduced RhoA signaling. On freshly isolated single myofibers, pMLC, a downstream target of active RhoA,^45^ was markedly decreased in MuSC-RhoA^fl/+^ compared to control MuSCs (Figure 2D, 2F), indicating reduced actomyosin contractility. FACS-isolated MuSCs displayed reduced F-actin intensity, visualized by Phalloidin staining (Figure 2G, 2I), consistent with diminished cytoskeletal tension. Aligned with these molecular changes, MuSC-RhoA^fl/+^ stem cells adopted a mechanically softened morphology characterized by reduced cell area and increased circularity (Figures 2H, 2J and 2K). This phenotype mirrored the low-tension state observed on soft substrates (Figures 1C, 1D), linking intrinsic RhoA activity to stiffness-dependent regulation of MuSC state. This shift was accompanied by an increased fraction of MuSCs expressing MyoD and the mechanosensitive transcriptional regulator YAP1 (Figures 2L - 282 cl:100 2O). These findings demonstrate that RhoA-dependent cytoskeletal tension is required to maintain the quiescent state of MuSCs *in vivo*.

To determine whether RhoA directly limits MuSC activation, we inhibited RhoA activity *in vitro* in WT MuSCs on isolated myofibers. Single muscle fibers were cultured in DMEM containing 10% horse serum media and treated with increasing concentrations of RhoA inhibitor (Rho1; 0, 0.5, or 1 µg/ml) for 8 hours (Supplementary Figure 1A). RhoA inhibition resulted in an increase in MyoD expression (Supplementary Figure 1B), recapitulating the *in vivo* genetic phenotype. Together, these results identify RhoA as an intrinsic regulator that prevents MuSC activation both *in vitro* and *in vivo*.

### RhoA maintains quiescence by inhibiting activation and cytoskeletal organization gene programs

To identify the transcriptional consequences of RhoA reduction, we performed RNA sequencing on freshly isolated MuSCs from control and MuSC-RhoA^fl/+^ mice. Differential gene expression analysis revealed that a greater number of genes were upregulated (347 genes) than downregulated (169 genes) (Supplementary Figure 2A). Among the upregulated transcripts, gene ontology (GO) terms associated with cell cycle progression^46^ were enriched, consistent with the premature cell cycle entry of the MuSCs at homeostasis.^15^ GO terms related to cytoskeletal remodeling and cell projection organization^47^ were also enriched, along with increased MAP kinase and integrin-mediated signaling (Supplementary Figure 2B).^12,48^ These processes are upregulated during the transition from quiescence to activation in MuSCs, indicating that RhoA downregulated MuSCs are shifting towards a more activated and mechanically responsive state.

In contrast, GO terms associated with cell-cell adhesion,^49^ including integrin-mediated adhesion and adherens junction maintenance, were downregulated, suggesting a loss of niche interactions. We also observed reduced expression of pathways involved in sensing mechanical stimuli and regulating calcium release from the sarcoplasmic reticulum,^50^ pointing to impaired mechanosensing. Additionally, downregulation of fatty acid oxidation is consistent with metabolic reprogramming that accompanies MuSC activation (Supplementary Figure 2C).^6,51^ Together, these data position RhoA as a central regulator of the transcriptional networks that coordinate cell cycle restraint, mechanical stability, and niche engagement in MuSCs.

### Gene body DNA methylation is associated with transcriptional and splicing changes in RhoA-deficient MuSCs

As DNA methylation is a central regulator of cell identity,^18,19^ we investigated whether RhoA-dependent MuSC activation was associated with alterations in DNA methylation pathways. Consistent with this possibility, RNA-seq transcripts per million (TPM) analysis revealed reduced Dnmt3a expression in MuSC-RhoA^fl/+^ stem cells compared to controls (data not shown). We therefore performed enzymatic methylation sequencing (EM-seq) on freshly isolated MuSCs from control and MuSC-RhoA^fl/+^ mice. This analysis identified 41,244 differentially methylated regions (DMRs; ≥10% methylation difference). DMRs were predominantly located within gene-associated regions (43%) and intergenic regions (39%) (Figures 3A). Of the gene-associated DMRs, the majority localized to gene bodies (89%), whereas only 8% mapped to promoter regions (Figure 3B), indicating that RhoA loss primarily alters DNA methylation within gene bodies rather than promoters.

**Figure 3.**
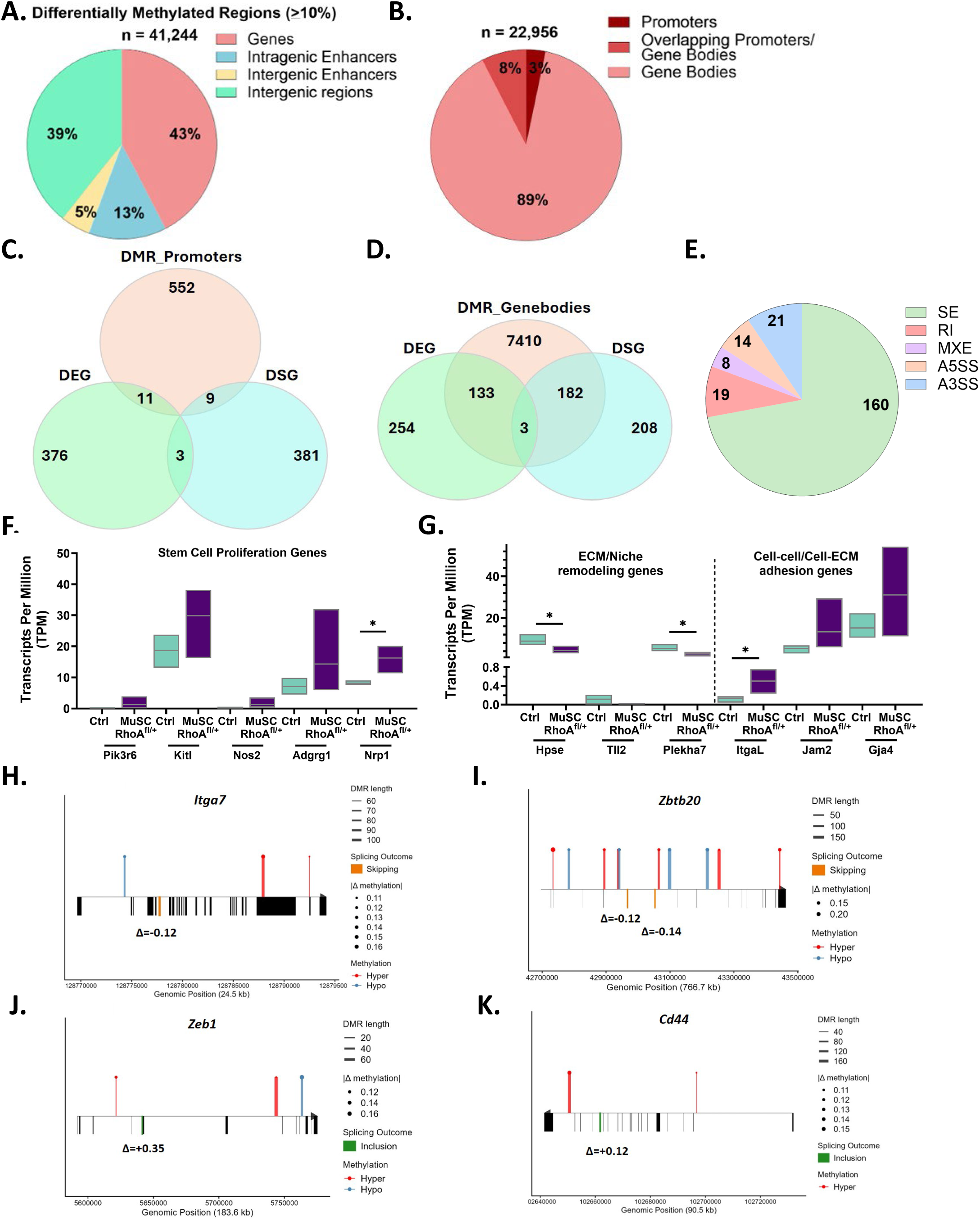
RhoA maintains the DNA methylation landscape in quiescent MuSCs. (A) Distribution of differentially methylated regions (DMRs) across genomic features (>10% methylation difference) in MuSCs freshly isolated from MuSC-RhoA^fl/+^ muscle compared to control. (B) Distribution of DMRs within gene regions. (C) Venn diagram showing overlap between promoter DMRs, differentially expressed genes (DEGs), and differentially spliced genes (DSGs). (D) Venn diagram showing overlap between gene body DMRs, DEGs, and DSGs. (E) Distribution of differentially spliced events in MuSC-RhoA^fl/+^ versus control MuSCs. (F) Representative stem cell proliferation genes containing both gene body DMRs and differential expression. (G) Representative niche remodeling and adhesion genes containing both gene body DMRs and differential expression. (H - K) Representative genes containing both gene body DMRs and alternative splicing events. (H, I) exons skipped upon RhoA reduction in MuSCs compared to control. (J, K) exons included upon RhoA reduction in MuSCs compared to control.

To define the functional impact of these changes, we integrated DMRs with transcriptional and splicing datasets. Promoter-associated DMRs showed minimal overlap with either differentially expressed genes (DEGs; 11 genes) or differentially spliced genes (DSGs; 9 genes) (Figure 3C). In contrast, gene body DMRs exhibited substantial overlap with both DEGs (136 genes) and DSGs (185 genes) (Figure 3D), suggesting that gene body methylation may serve as an interface linking DNA methylation to transcriptional output and splice isoform diversity. Differential splicing events spanned multiple categories, including skipped exons (SE), retained introns (RI), mutually exclusive exons (MXE), alternative 5ʹ splice sites (A5SS), and alternative 3ʹ splice sites (A3SS), with skipped exon events representing the predominant class of splicing alterations (Figure 3E).

Consistent with previous findings, gene body methylation was largely associated with gene activation.^22–24^ Among the 136 overlapping genes between DMR in gene body and DEG, 112 were upregulated and 24 were downregulated, despite a balanced distribution of hyper- and hypomethylated DMRs (Supplementary Figure 2D, 2E). GO analysis revealed enrichment in cell-matrix adhesion, focal adhesion, and stem cell differentiation, indicating activation of programs that promote niche remodeling and exit from quiescence (Supplementary Figure 3F). Analysis of transcripts per million (TPM) values for genes with both gene body DMRs and DEGs further supported this transition. Genes linked to proliferation in MuSCs or other stem cell systems (e.g., Pik3r6, Kitl, Nos2, Adgrg1, and Nrp1)^52–56^ were upregulated in RhoA-deficient cells (Figure 3F), alongside increased expression of differentiation-associated genes (Dysf, Myh7b, Tbx5, Casz1, and Mitf)^57–61^ (Supplementary Figure 2G). In contrast, genes involved in extracellular matrix remodeling (Hpse, Tll2, and Plekha7)^62–64^ were down regulated, while adhesion-related genes (ItgaL, Jam2, and Gja4)^65–67^ were elevated (Figure 3G), collectively indicating a shift toward activation accompanied by altered niche interactions.

Genes exhibiting both gene body methylation changes and alternative splicing were enriched for pathways regulating cell differentiation, glycolysis, morphogenesis, and adhesion, along with chromatin remodeling and epigenetic regulation (Supplementary Figure 2H), supporting a coordinated role for gene body methylation in coupling transcriptional control with splice isoform selection. Specific splicing events further illustrate this relationship between DMR in gene bodies and DSG. *Itga7*, a key laminin receptor in MuSCs, undergoes alternative splicing of mutually exclusive exons encoding extracellular domains that determine laminin binding specificity.^65^ RhoA-deficient MuSCs preferentially skipped exon 6, preceded by a hypomethylated DMR (Figure 3H), while *Zbtb20*, a regulator of MuSC establishment and lineage progression,^68^ exhibited skipping of two adjacent exons associated with hypomethylation (Figure 3I), highlighting a link between reduced methylation and exon exclusion in genes governing MuSC identity. Conversely, exon inclusion events were associated with hypermethylation. *Zeb1*, which is required for MuSC regeneration following injury,^69^ showed increased exon inclusion downstream of a hypermethylated DMR (Figure 3J), and *Cd44*, a hyaluronic acid receptor involved in MuSC activation and migration,^70,71^ displayed a similar pattern (Figure 3K), indicating that increased methylation is linked to exon inclusion in genes regulating stem cell state transitions.

Collectively, these findings demonstrate that disruption of RhoA-dependent mechanical signaling reshapes gene body methylation to coordinate transcriptional and splicing programs, thereby promoting exit from quiescence and altering MuSC interactions with the niche.

### RhoA-dependent mechanical signaling regulates the expression of Dnmt3a in quiescent MuSCs

To investigate whether mechanical signaling regulates DNA methylation machinery, we examined enzymes responsible for the establishment and maintenance of CpG methylation. Immunofluorescence analysis of MuSCs on freshly isolated myofibers revealed that Dnmt1 expression was low in quiescent MuSCs and remained unchanged upon reduction of RhoA signaling (Figures 4A, 4B). Dnmt3B was highly expressed in quiescent MuSCs but was not affected by reduced RhoA signaling (Figures 4F, 4G). In contrast, Dnmt3A protein levels were significantly reduced in MuSC-RhoA^fl/+^ stem cells relative to controls (Figures 4C, 4D). Consistent with these findings, qRT-PCR analysis on FACS-isolated MuSCs showed a significant decrease in Dnmt3a transcript levels following the loss of RhoA signaling (Figure 4E), identifying Dnmt3a as a downstream target of the RhoA-dependent mechanical signaling pathway. To further test this relationship, pharmacological inhibition of RhoA on isolated WT myofibers similarly reduced Dnmt3A expression (Supplementary Figure 3A), recapitulating the *in vivo* genetic phenotype and demonstrating that Dnmt3A expression is dynamically regulated by RhoA activity.

**Figure 4.**
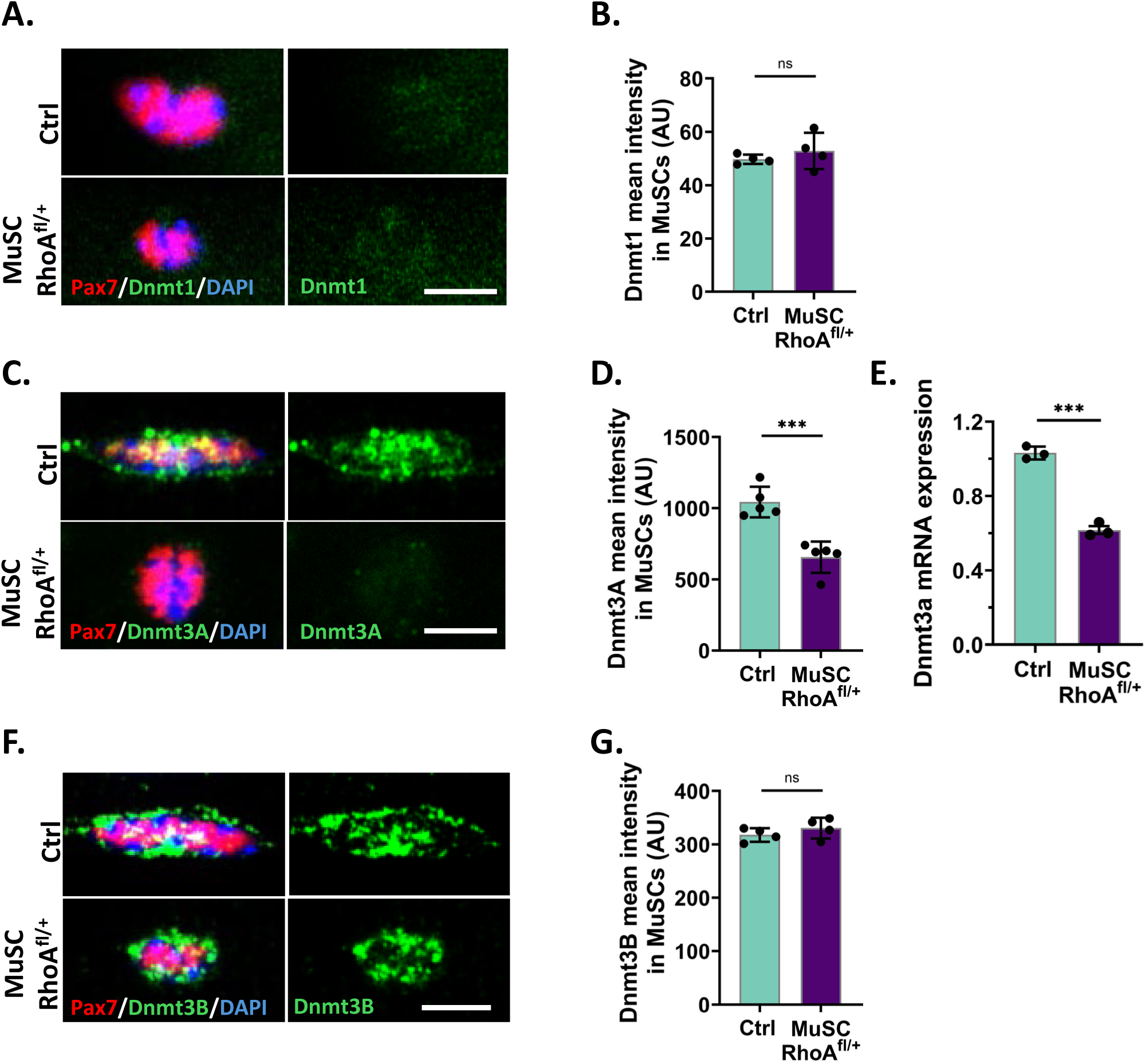
RhoA selectively regulates Dnmt3A expression in quiescent MuSCs. (A and B) Representative images (A) and quantification (B) of Dnmt1 intensity in MuSCs on control and MuSC-RhoA^fl/+^ myofibers (n = 4). (C and D) Representative images (C) and quantification (D) of Dnmt3A intensity in MuSCs on control and MuSC-RhoA^fl/+^ myofibers (n = 5). (E)) Dnmt3a mRNA expression in freshly isolated MuSCs from control and MuSC-RhoA^fl/+^ mice 30 days after tamoxifen injection. Expression levels were normalized to GAPDH, and fold change relative to control samples was determined using the ΔΔCt method (n = 3). (F and G) Representative images (F) and quantification (G) of Dnmt3B intensity in MuSCs on control and MuSC-RhoA^fl/+^ myofibers (n = 4). Each dot in bar graphs represents an individual mouse. Statistical significance was determined using an unpaired two-tailed Student’s t test. Error bars, mean ± SD; ***p < 0.001, ns- not significant; scale bars, 5µm in (A, C, and F).

We next asked whether extracellular stiffness regulates Dnmt3A expression. MuSCs cultured on matrices of defined stiffness (0.5, 16, and 32kPa) displayed a stiffness-dependent increase in Dnmt3A levels with reduced levels under soft conditions that mimic reduced RhoA signaling (Supplementary Figure 3B). Thus, extrinsic mechanical cues and intrinsic RhoA activity together regulate the expression of Dnmt3A.

### Dnmt3A Maintains the Quiescent State of MuSCs

We focused our subsequent analyses on Dnmt3A due to its role as a *de novo* DNA methyltransferase and its established functions in stem cell fate regulation.^26,72^ To determine whether Dnmt3A maintains MuSC quiescence, we reduced the levels of *Dnmt3a* in adult MuSCs using Pax7Cre^ER^; Dnmt3a^fl/+^ mice.^43,73^ These mice will be referred to as MuSC-Dnmt3A^fl/+^, with Cre negative Dnmt3a^fl/+^ littermates as controls (Ctrls). Tamoxifen administration for 7 consecutive days followed by a 30-day recovery period resulted in a reduction in Dnmt3A protein levels in MuSCs (Figures 5A-5C). MuSC activation was measured by immunostaining for MyoD (Figures 5D, 5E) and YAP1 (Figures 5F, 5G) expression, which were significantly increased in MuSC-Dnmt3A^fl/+^ cells compared with controls. These findings indicate that Dnmt3A is required to maintain MuSC quiescence, and that reduced Dnmt3A levels are sufficient to drive a spontaneous transition toward MuSC activation.

**Figure 5.**
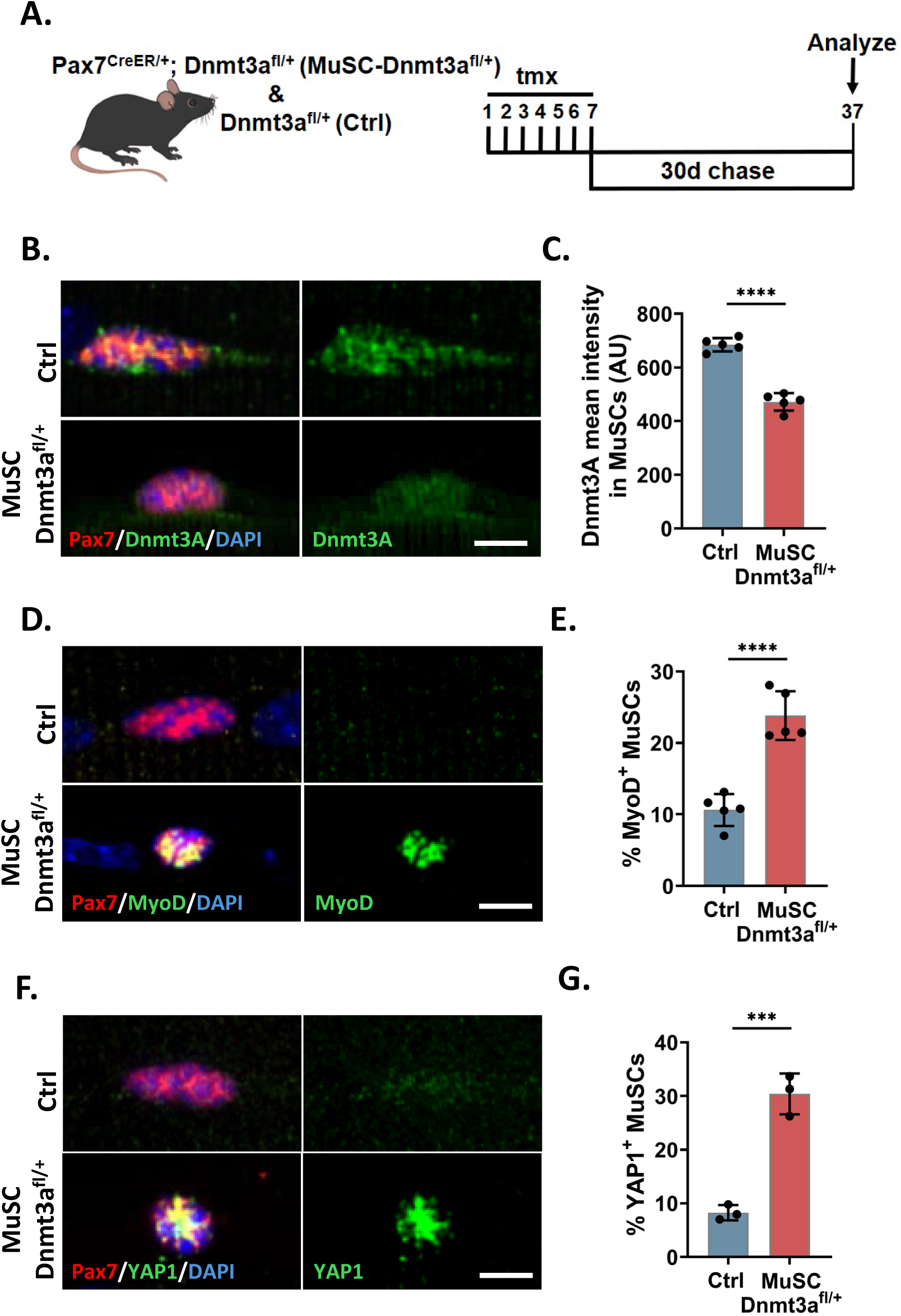
Dnmt3A Maintains MuSC Quiescence. (A) Schematic of the experimental design (B and C) Representative images (A) and quantification (B) of Dnmt3A intensity in MuSCs on control and MuSC-Dnmt3A^fl/+^ myofibers (n = 5). (D and E) Representative images (D) and quantification (E) of the percent of MyoD positive MuSCs on control and MuSC-Dnmt3A^fl/+^ myofibers (n = 5). (F and G) Representative images (F) and quantification (G) of the percent of YAP1 positive MuSCs on control and MuSC-Dnmt3A^fl/+^ myofibers (n = 3). Each dot in bar graphs represents an individual mouse. Statistical significance was determined using an unpaired two-tailed Student’s t test. Error bars, mean ± SD; ***p < 0.001, ****p < 0.0001; scale bars, 5µm in (B, D, and F).

We next examined whether loss of *Dnmt3a* alters cytoskeletal tension. pMLC levels were significantly reduced in MuSC-Dnmt3A^fl/+^ stem cells (Supplementary Figures 4A, 4B), indicating diminished actomyosin contractility. Consistent with this, Dnmt3A-deficient MuSCs displayed a smaller cell area and increased circularity (Supplementary Figures 4C, 4D), phenocopying the low-tension, activation-prone state observed following reduction of RhoA signaling. These findings suggest that Dnmt3A not only functions downstream of RhoA signaling but also contributes to the maintenance of cytoskeletal tension and cell morphology associated with MuSC quiescence. Together, these data support a feedback relationship in which mechanical signaling through RhoA regulates Dnmt3A expression, while Dnmt3A reinforces the cellular architecture and tension state required to maintain stem cell quiescence.

### RhoA regulates the expression of Dnmt3A through SP1-dependent transcription

To define how RhoA signaling regulates *Dnmt3a* expression, we investigated the transcriptional mechanisms controlling *Dnmt3a* regulation. Specificity Protein 1 (SP1), a ubiquitously expressed zinc-finger transcription factor, is known to regulate both human *DNMT3A* and *DNMT3B* expression through binding to their GC-rich promoter regions.^74,75^ Immunostaining analysis revealed that total SP1 levels were unchanged upon reduction of RhoA signaling (Figures 6A, 6B). However, phosphorylation of SP1 at Thr453, a modification that reduces its DNA-binding activity in certain cellular contexts,^76^ was significantly increased in RhoA-knocked down MuSCs (Figures 6C, 6D), suggesting reduced SP1 transcriptional function.

**Figure 6.**
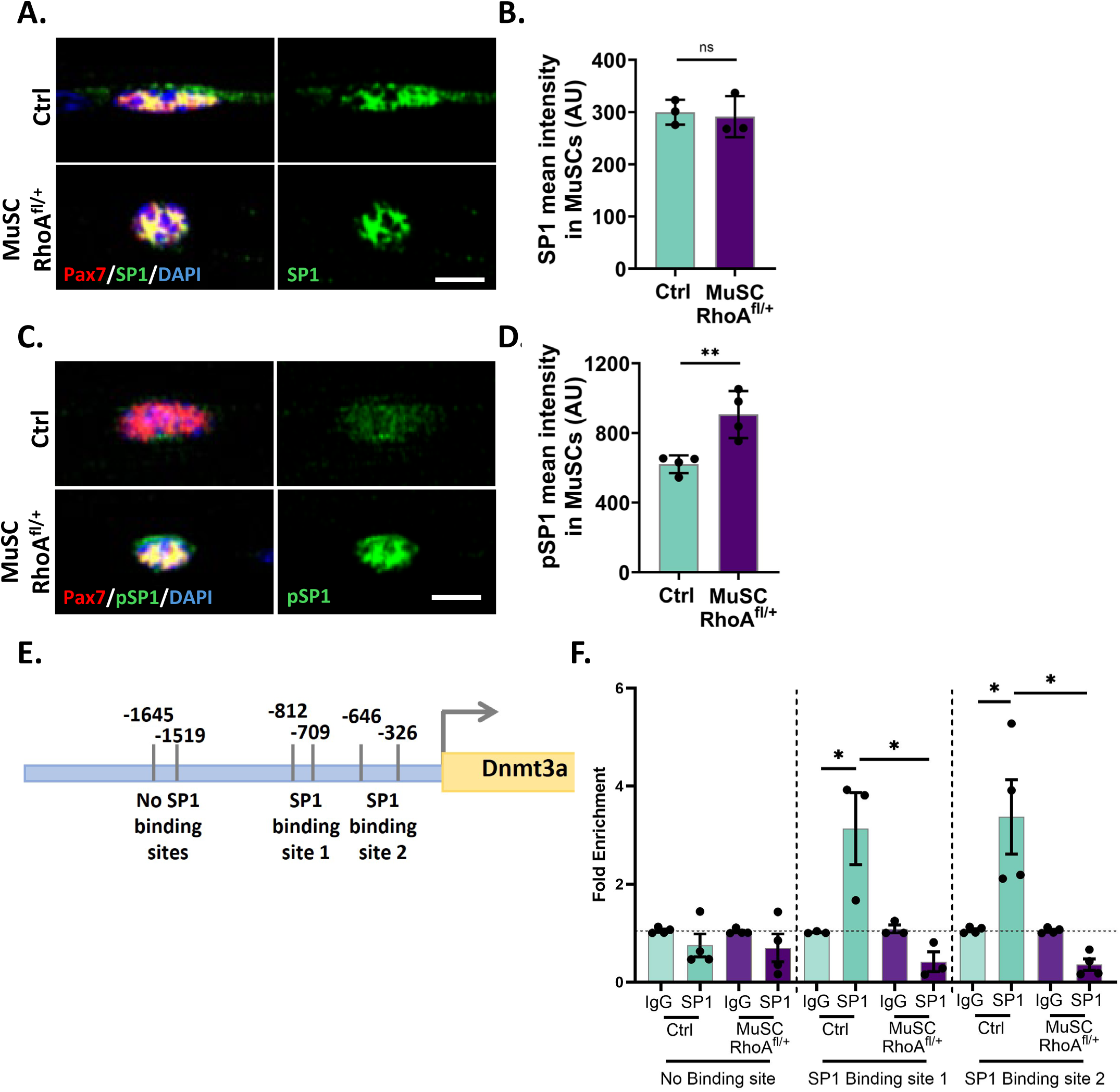
RhoA regulates Dnmt3A expression by modulating SP1 phosphorylation. (A and B) Representative images (A) and quantification (B) of SP1 intensity in MuSCs on control and MuSC-RhoA^fl/+^ myofibers (n = 3). (C and D) Representative images (C) and quantification (D) of phosphorylated SP1 (pSP1) intensity in MuSCs on control and MuSC-RhoA^fl/+^ myofibers (n = 4). (E) Schematic of SP1 binding sites on *Dnmt3a* promoter. (F) ChIP-qPCR analysis of SP1 occupancy at SP1-binding and non-binding regions of the *Dnmt3a* promoter in freshly isolated MuSCs from control and MuSC-RhoA^fl/+^ mice. SP1 ChIP signal was normalized to IgG control, and fold enrichment was determined using the ΔΔCt method (n≥3). Each dot in bar graphs represents an individual mouse. Statistical significance was determined using an unpaired two-tailed Student’s t test. Error bars, mean ± SD; *<0.05, **p < 0.01, ns- not significant; scale bars, 5µm in (A) and (C).

To test whether RhoA regulates SP1 occupancy at the *Dnmt3a* promoter, we performed Chromatin Immunoprecipitation followed by PCR (ChIP-PCR) for SP1 binding sites. SP1 occupancy at the *Dnmt3a* promoter was markedly reduced in MuSC-RhoA^fl/+^ stem cells relative to controls which displayed approximately three-fold higher SP1 enrichment at the promoter. (Figures 6E, 6F). These findings indicate that RhoA-dependent cytoskeletal tension promotes SP1-mediated binding, thereby sustaining Dnmt3a transcription and maintaining SC quiescence.

### SP1 activity is required to maintain Dnmt3A expression and SC quiescence *in vivo* and *in vitro*

To determine whether SP1 activity is required *in vivo*, WT mice were injected intraperitoneally with the SP1 inhibitor (Mitramycin A; 0, 0.5, 1 mg/kg) for 14 consecutive days (Figure 7A). SP1 inhibition led to reduced cell area and increased circularity (Figures 7B, 7C), indicating a mechanically softened state of the MuSCs at homeostasis. This was accompanied by increased MyoD expression (Figure 7D) and a marked reduction in Dnmt3A levels (Figure 7E), demonstrating that SP1 activity is required to maintain Dnmt3A expression and MuSC quiescence. Consistent with the known mechanism of Mithramycin A-mediated inhibition of SP1 DNA binding, total and phosphorylated SP1 levels remained unchanged (Figures 7F, 7G), indicating that the observed effects are independent of changes in SP1 abundance or phosphorylation.^77,78^ Body weight of mice and cleaved caspase-3 levels were unaffected (Supplementary Figures 5F, 5G), excluding systemic toxicity or apoptosis as confounding factors.

**Figure 7.**
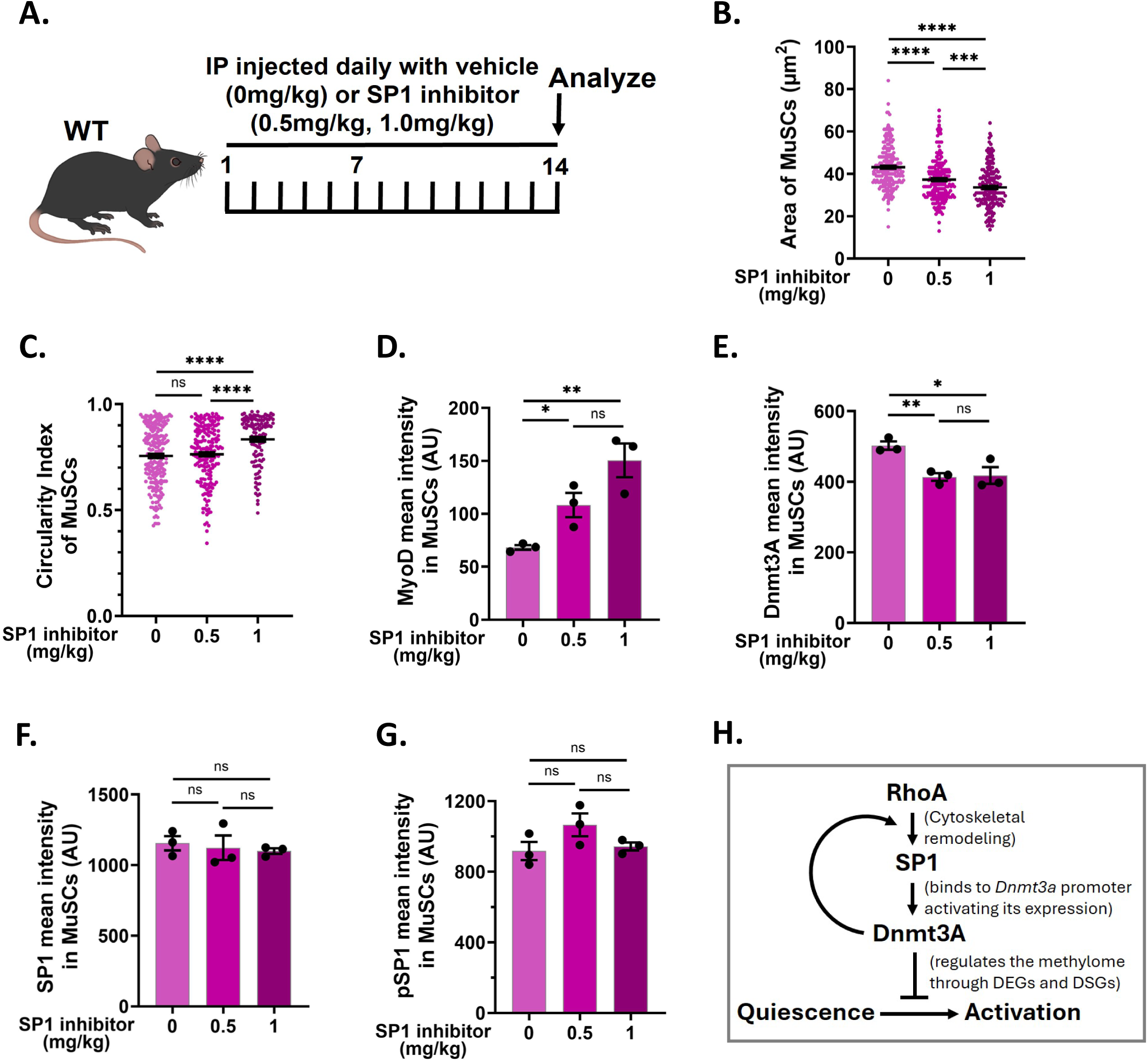
Pharmacological inhibition of SP1 *in vivo* promotes MuSC activation at homeostasis. (A) Schematic of the experimental design. (B - G) Quantification of cell area (B), circularity index (C), MyoD intensity (D), Dnmt3a intensity (E), SP1 intensity (F), and pSP1 intensity (G) in WT MuSCs on isolated myofibers from vehicle and SP1 inhibitor (Mitramycin A) treated mice at indicated concentrations for 14 consecutive days (n=3). (H) Model summarizing the proposed mechanism. Each dot in bar graphs represents an individual mouse; each dot in dot plots represents an individual MuSC. Statistical significance was determined using an unpaired two-tailed Student’s t test. For the bar graph, Error bars, mean ± SD; For the dot plots, Error bars, mean ± SEM *p<0.05, **p < 0.01, ***p < 0.001, ****p < 0.0001, ns- non significant.

To test whether SP1 directly regulates Dnmt3A in MuSCs, isolated WT myofibers were treated with Mithramycin A for 8 hours (Supplementary Figure 5A). Acute SP1 inhibition produced an increase in MyoD expression and a corresponding decrease in Dnmt3A (Supplementary Figures 5B, 5C), without altering total or phosphorylated SP1 levels (Supplementary Figures 5D, 5E). These findings demonstrate that SP1 DNA binding is required to sustain Dnmt3A expression and prevent premature MuSC activation.

Together, these findings identify a mechanosensitive epigenetic circuit that preserves MuSC quiescence. Physiological stiffness establishes a quiescent state that is maintained by RhoA-dependent cytoskeletal tension, which sustains SP1-mediated transcription of Dnmt3a. Dnmt3A protein, in turn, reinforces cytoskeletal tension to stabilize this state and prevent premature activation at homeostasis. This RhoA-SP1-Dnmt3A axis couples mechanical inputs from the niche to epigenetic regulation of stem cell fate (Figure 7H).

## DISCUSSION

Stem cell function is influenced by signals from the niche that integrate biochemical and mechanical inputs to regulate quiescence, activation, and self-renewal.^79^ While the role of mechanical cues in controlling stem cell behavior is increasingly appreciated,^37,80^ how these signals are translated into stable molecular programs that preserve stem cell identity remains unclear. Here, we define a mechanosensitive epigenetic mechanism by which physiological muscle stiffness maintains MuSC quiescence. Our findings show that RhoA-dependent cytoskeletal tension maintains the DNA methylation landscape through regulation of *Dnmt3a*, thereby coordinating transcriptional and alternative splicing programs that sustain the quiescent state.

Mechanical signals from the niche have been shown to influence stem cell fate decisions through effects on cytoskeletal organization and downstream signaling pathways.^80,81^ However, these inputs are often considered transient and reversible. Our data suggest that mechanical cues can be encoded into the epigenome, providing a mechanism for stabilizing cell state. Notably, we find that mechanical signaling regulates the expression of a core DNA methylation enzyme, Dnmt3A, linking cytoskeletal tension directly to the establishment of the epigenetic landscape. These findings support a model in which physical properties of the niche are not only sensed but are translated into durable molecular states that reinforce stem cell identity.

Among the epigenetic regulators examined, Dnmt3A emerged as a key regulator of this process. Dnmt3A is a *de novo* DNA methyltransferase with established roles in stem cell function across multiple tissues,^26,82,83^ yet its integration with mechanical signaling has not been defined. We show that RhoA activity sustains Dnmt3a expression, positioning it as a downstream effector of cytoskeletal tension. Mechanistically, we identify SP1 as a transcriptional regulator linking RhoA signaling to Dnmt3a expression. RhoA-dependent cytoskeletal tension promotes SP1 occupancy at the *Dnmt3a* promoter, and inhibition of SP1 *in vivo* is sufficient to drive MuSC activation under homeostatic conditions. These findings position SP1 as a key intermediary through which mechanical signals are translated into transcriptional control of the epigenetic machinery.

While Dnmt3A emerged as a central mechanosensitive regulator, our data also suggest that epigenetic remodeling in MuSCs reflects the combined activity of multiple DNA methylation and demethylation enzymes. Notably, Dnmt3B expression remained unchanged despite alterations in mechanical signaling, suggesting that global methylation capacity may be maintained even as Dnmt3A levels are modulated.^84^ This may explain why both hypermethylation and hypomethylation events are observed, rather than a unidirectional shift in the methylome. These findings support a model in which RhoA-dependent mechanical signaling selectively modulates epigenetic regulators to preserve stem cell identity and quiescence.

In addition to transcriptional regulation, our data support a role for DNA methylation in controlling alternative splicing in MuSCs. Although DNA methylation has been primarily studied in the context of gene expression, emerging evidence indicates that it can influence splice site selection and isoform diversity.^33,83^ Our findings extend this concept to adult stem cells and suggest that RhoA-dependent maintenance of the DNA methylome contributes to the regulation of both transcriptional output and splicing programs. This coupling provides a mechanism by which a single epigenetic layer can coordinate multiple aspects of stem cell identity, including gene expression and isoform diversity.

Together, these findings support a model in which MuSC quiescence is not a passive default state, but rather an actively maintained condition reinforced by mechanical cues from the niche.^85^ Physiological stiffness establishes a quiescent mechanical state that is sustained by RhoA-dependent cytoskeletal tension, which in turn preserves the epigenetic landscape required for stem cell identity. Disruption of this pathway, through loss of RhoA signaling, leads to remodeling of the DNA methylome and shifts in transcriptional and splicing programs consistent with activation.

While our study identifies Dnmt3A as a central mediator of mechanosensitive epigenetic regulation, several questions remain. The specific genomic loci at which DNA methylation changes drive transcriptional versus splicing outcomes have yet to be defined. In addition, the extent to which Dnmt3A cooperates with other methyltransferases and TET enzymes to regulate these processes will require further investigation. Future studies will also be needed to determine how mechanical cues intersect with other signaling pathways in the niche to fine-tune stem cell behavior.

More broadly, these findings have implications for understanding how changes in tissue mechanics during aging, injury, or disease may alter stem cell function. Alterations in muscle stiffness are a feature of both aging and pathological conditions,^86^ and our results suggest that such changes could impact MuSCs behavior through epigenetic mechanisms. Defining how mechanical signals are integrated into the epigenome may provide new strategies to modulate stem cell function in regenerative and disease contexts.

In summary, we define a mechanical-epigenetic circuit that converts niche-derived physical cues into a stable molecular program that preserves muscle stem cell quiescence. These findings establish a framework linking cytoskeletal tension to DNA methylation and provide insight into how mechanical dysregulation may contribute to impaired tissue regeneration.

---

**Supplementary Figure 1.**
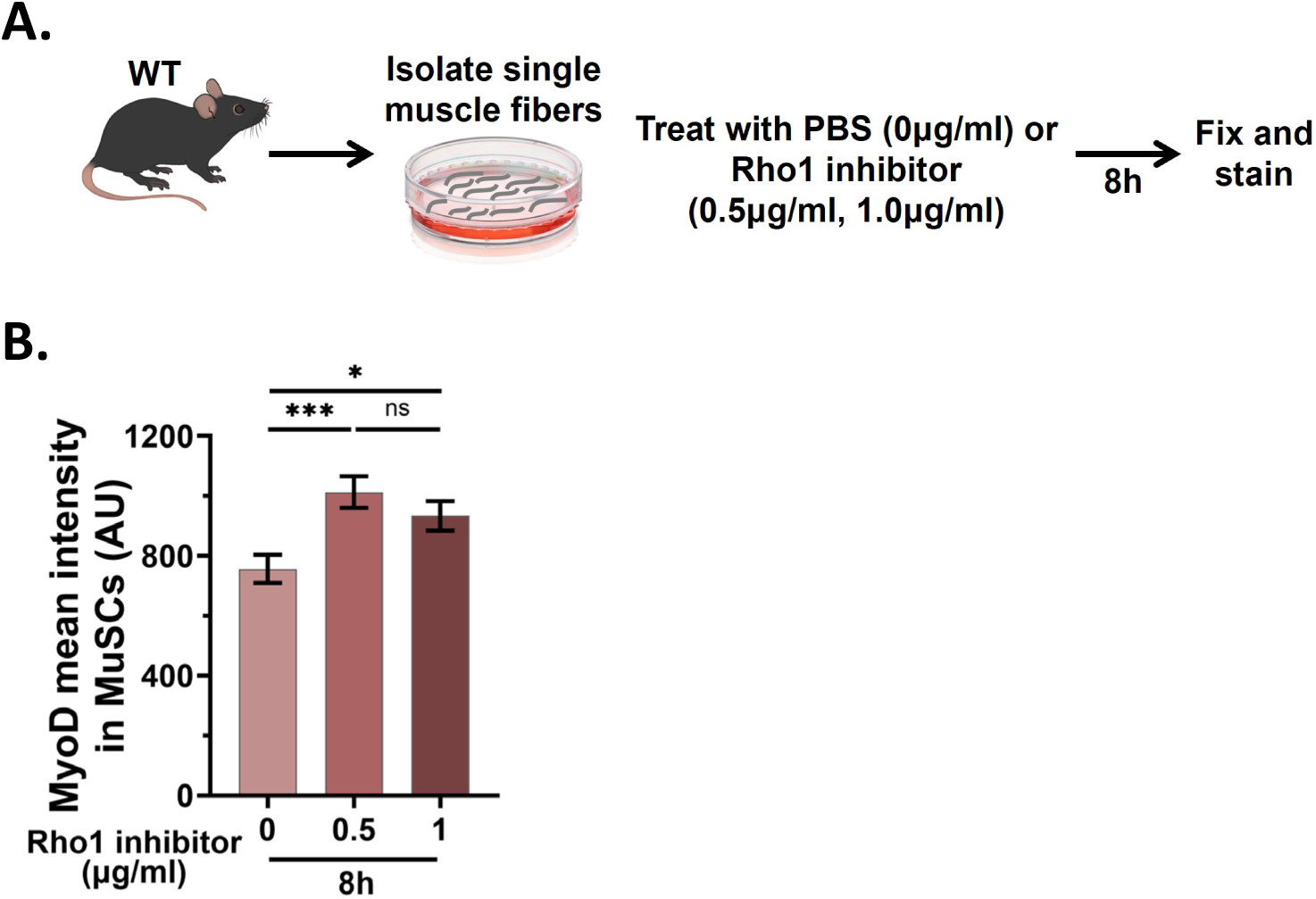
Inhibition of RhoA *in vitro* accelerates MuSC activation Related to Figure 2. (A) Schematic of the experimental design. (B) Quantification of MyoD intensity in MuSCs on isolated WT myofibers that were treated immediately after isolation with increasing concentrations of a RhoA inhibitor (Rho1) for 8 hours (n = 4).

**Supplemental Figure 2.**
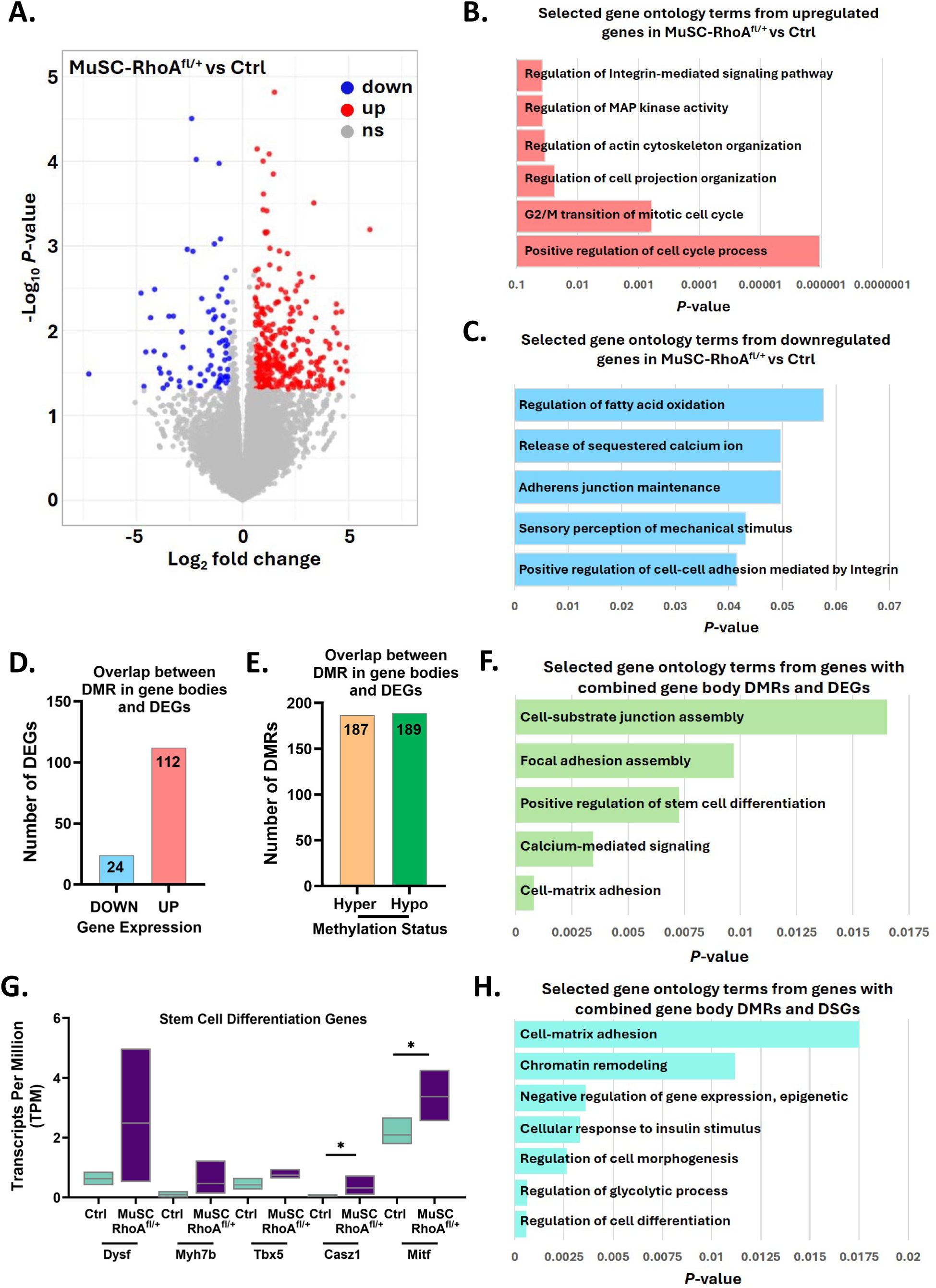
RhoA-dependent transcriptional and epigenetic programs define MuSC state Related to Figure 3. (A) Volcano plot of RNA-seq data showing upregulated and downregulated genes in freshly isolated MuSCs from MuSC-RhoA^fl/+^ muscle compared to control. (B and C) Gene Ontology (GO) analysis of upregulated (B) and downregulated (C) genes. (D) Distribution of DEGs overlapping with gene body DMRs, categorized as upregulated or downregulated. (E) Distribution of gene body DMRs overlapping with DEGs, categorized as hypermethylated or hypomethylated. (F) GO analysis of genes containing both gene body DMRs and differential expression. (G) Representative stem cell differentiation genes containing both gene body DMRs and differential expression. (H) GO analysis of genes containing both gene body DMRs and DSGs.

**Supplementary Figure 3.**
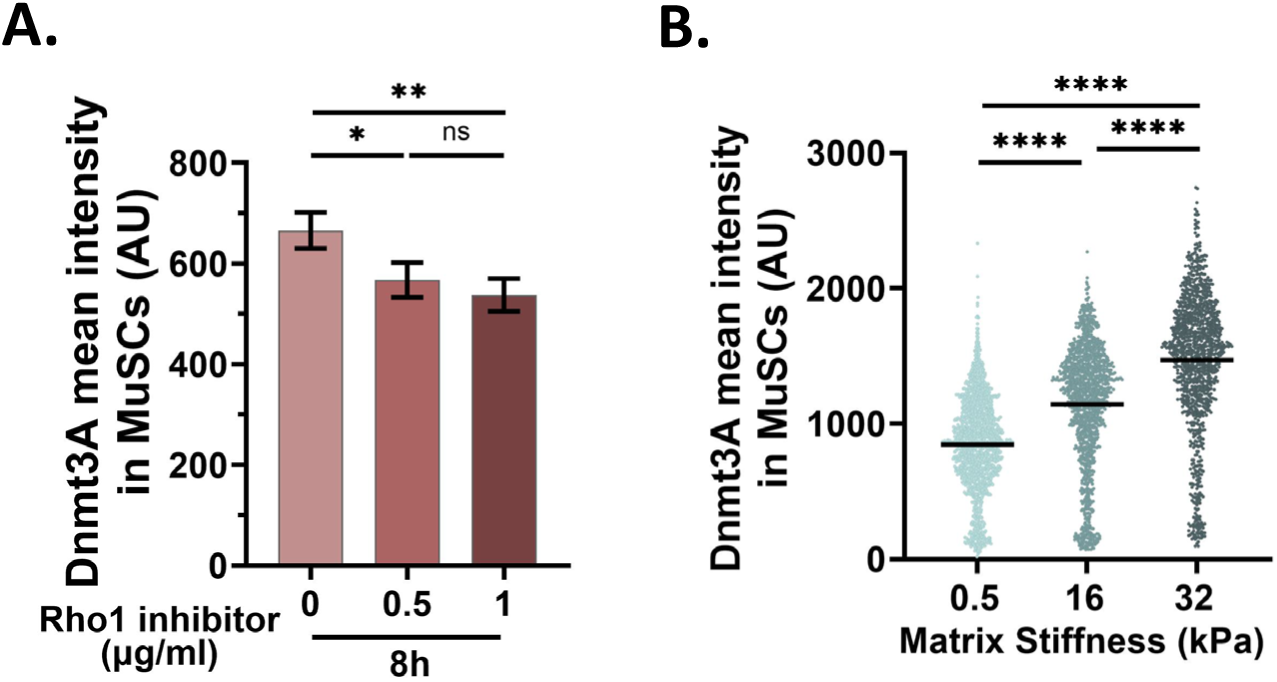
RhoA-dependent mechanical signaling regulates Dnmt3A expression in MuSCs Related to Figure 4. (A) Quantification of Dnmt3A intensity in WT MuSCs treated immediately after isolation with increasing concentrations of a RhoA inhibitor for 8 hours (n = 4). (B) Quantification of Dnmt3A intensity in SCs plated on matrices of defined stiffness (0.5, 16 and 32 kPa) for 10 hours (n=4). Each dot in bar graphs represents an individual mouse; each dot in dot plots represents an individual MuSC, with the central line indicating the median. Statistical significance was determined using an unpaired two-tailed Student’s t test. Error bars, mean ± SD; *p<0.05, **p < 0.01, ****p < 0.0001, ns- not significant.

**Supplementary Figure 4.**
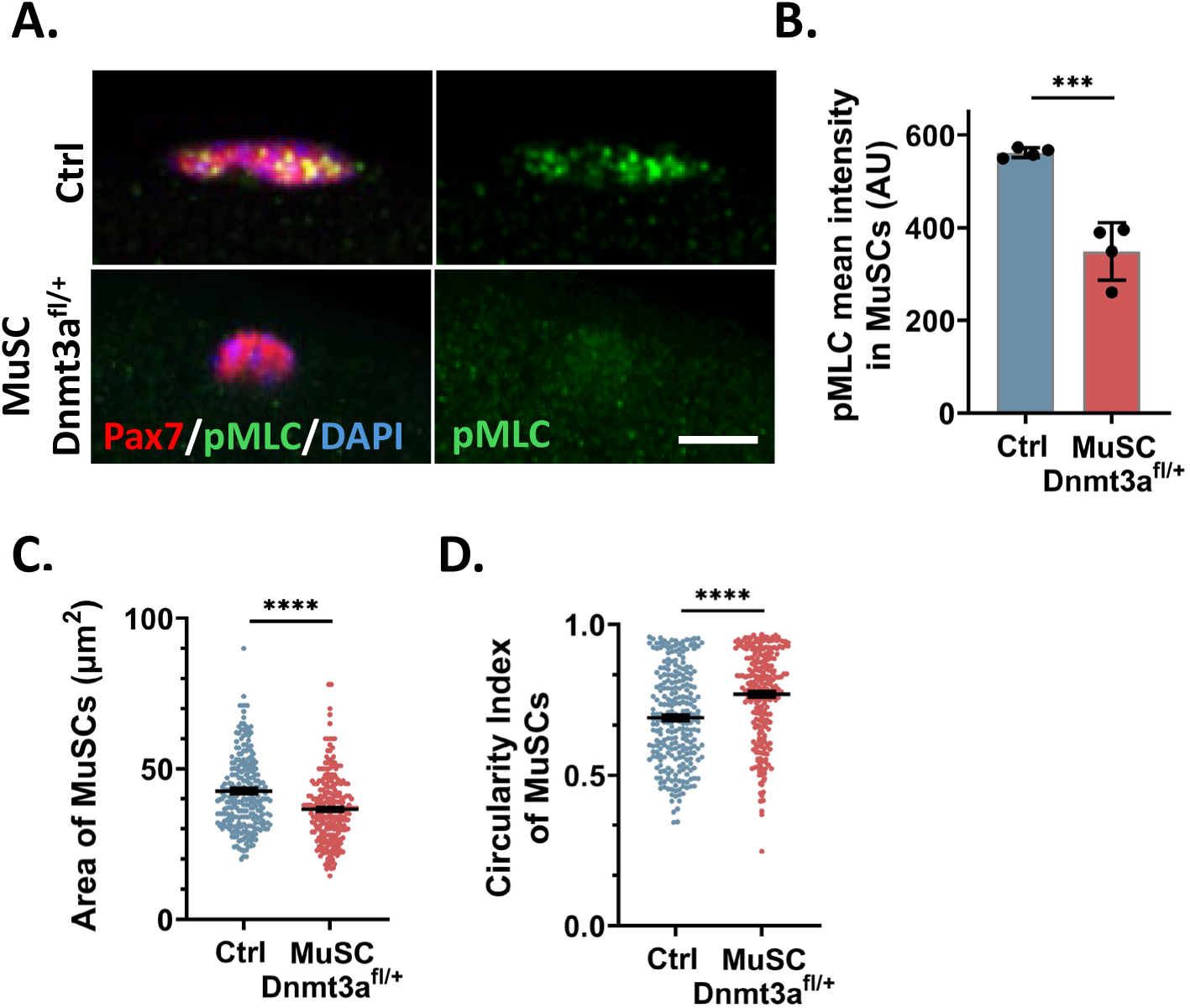
Dnmt3A loss phenocopies RhoA depletion in MuSCs Related to Figure 5. (A and B) Representative images (A) and quantification (B) of pMLC intensity in MuSCs on control and MuSC-Dnmt3A^fl/+^ myofibers (n = 4). (C and D) Quantification of cell area (C) and circularity (D) of MuSCs on control and MuSC- Dnmt3a^fl/+^ myofibers. Each dot in bar graphs represents an individual mouse; each dot in dot plots represents an individual MuSC. Statistical significance was determined using an unpaired two-tailed Student’s t test. For the bar graph, Error bars, mean ± SD; For the dot plots, Error bars, mean ± SEM; ***p < 0.001, ****p < 0.0001; scale bars, 5µm in (A).

**Supplementary Figure 5.**
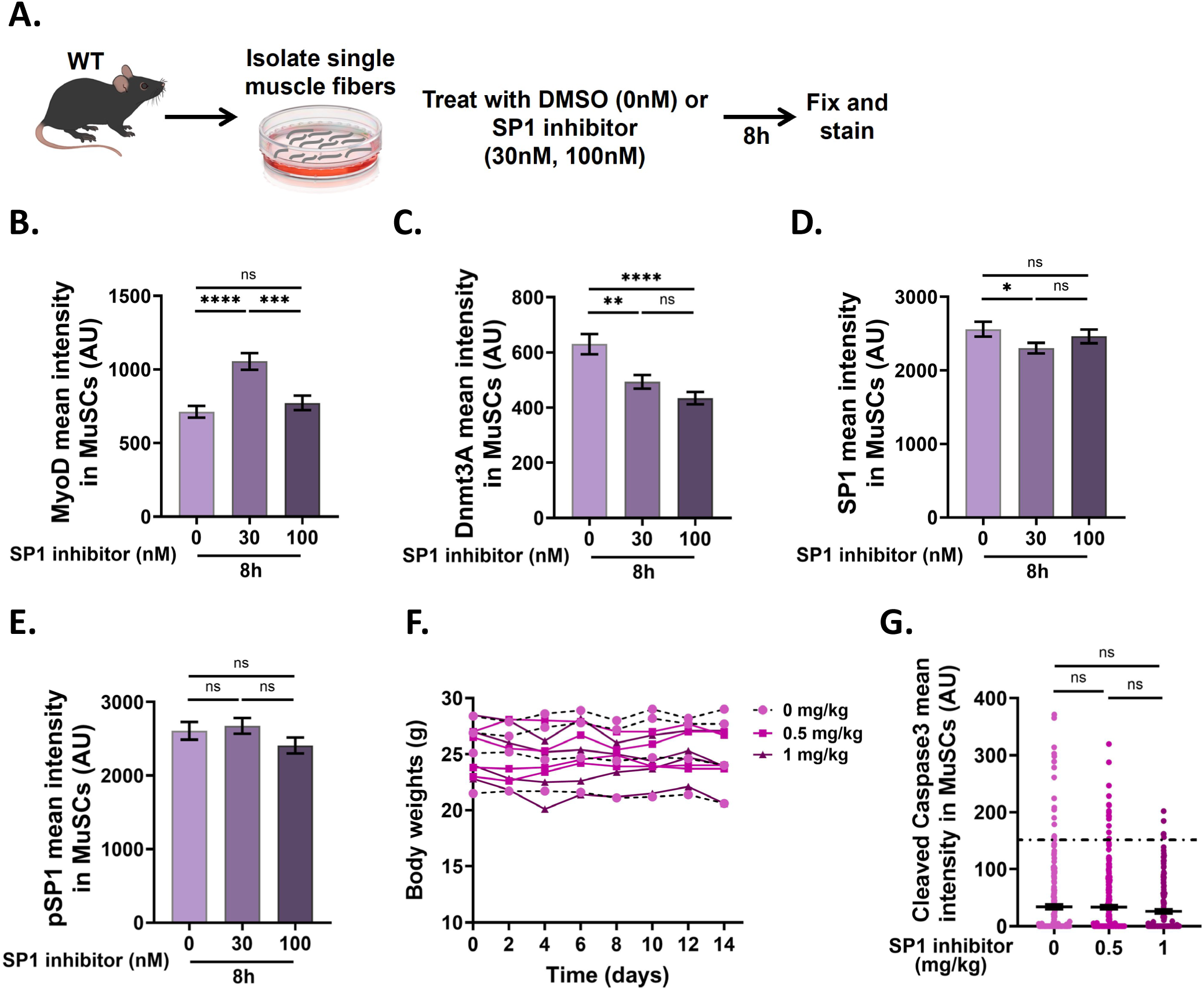
SP1 Inhibition *in vitro* promotes MuSC activation and reduces Dnmt3A expression Related to Figure 7. (A) Schematic of the experimental design. (B - E) Quantification of MyoD intensity (B), Dnmt3A intensity (C), SP1 intensity (D) and pSP1 intensity in WT MuSCs on isolated myofibers treated with vehicle or SP1 inhibitor *in vitro* for 8 hours (n = 3). (F) Body weight of mice treated with indicated doses (0, 0.5 and 1mg/kg) of SP1 inhibitor for 14 consecutive days. Each line represents an individual mouse measured every other day (n = 3). (G) Quantification of cleaved caspase 3 intensity in MuSCs on isolated myofibers from vehicle and SP1 inhibitor treated mice. Each dot in the dot plot represents an individual MuSC, with the error bar, mean ± SEM (n = 3). Statistical significance was determined using an unpaired two-tailed Student’s t test. For the bar graphs, Error bars, mean ± SD; *p<0.05, **p < 0.01, ***p < 0.001, ****p < 0.0001, ns- non significant.

## METHODS

### Experimental Model

Mice used in this study were housed and maintained at the University of North Dakota (UND) Center for Biomedical Research (CBR). All animal experiments were approved by the Institutional Animal Care and Use Committee (IACUC) and performed in compliance with applicable federal and state guidelines and regulations. Previously published *Pax7Cre^ER^*,^43^ *RhoA^flox/+^*,^44^ and *Dnmt3a^flox/+^* ^73^ mouse lines were used in this study. Wild-type (WT) C57BL/6 mice were obtained from the Jackson Laboratory. All mice used were adults between the age of 3 - 5 months. Control and experimental animals were littermates for all experiments, and equal numbers of male and female mice were used. Animals were genotyped by Polymerase Chain Reaction (PCR) using DNA isolated from toe clips collected from pups younger than 7 days of age. Genotyping primer sequences are available upon request.

### Method Details

#### Animal procedures

Tamoxifen (Sigma-Aldrich) was prepared in corn oil at a concentration of 20 mg/mL. To induce Cre-mediated gene deletion, control and experimental mice received intraperitoneal injections of tamoxifen at a dose of 150 mg/kg/day for seven consecutive days. Mice were then allowed to recover for one month prior to MuSC analysis.

To determine the role of SP1 in maintaining MuSC quiescence, WT mice were treated with Mitramycin A (Cayman Chemical), an inhibitor of SP1 activity. Mice were assigned to treatment groups receiving 0, 0.5, or 1 mg/kg Mitramycin A. Body weight was recorded prior to the initial injection and monitored every other day throughout the 14-day treatment period to assess changes in animal health and weight. Mitramycin A (Cayman Chemical) was initially prepared in sterile DMSO and further diluted in 1× PBS to achieve the desired dosing concentrations. Mice received daily intraperitoneal injections at a volume of 150-200 µL per mouse for 14 consecutive days.

### FACS isolation of muscle stem cells

Forelimb, hindlimb, and back muscles were dissected from mice, finely minced, and digested in DMEM containing 0.2% Collagenase Type 2 (Worthington) in a shaking water bath at 37°C for 90 minutes. Muscle tissue was then washed and subjected to a second digestion with 0.2% Collagenase (Worthington) and 0.4% Dispase (Gibco) in a shaking water bath at 37°C for 30 minutes. The digested tissue was passed through a 20-gauge needle three times, then filtered sequentially through 40 μm and 30 μm strainers to obtain a single cell suspension. Cells were incubated with fluorophore conjugated antibodies: PE-Cy7 anti-mouse CD31 (clone: 390; BD Biosciences), PE-Cy7 anti-mouse CD45 (clone: 30-F11; BD Biosciences), APC-Cy7 anti-mouse Sca1 (clone: D7; BD Biosciences), PE anti-mouse CD106/VCAM-1 (clone: M/K-2; Invitrogen) and APC anti-mouse α7 integrin (clone: R2F2; AbLab), for one hour at room temperature. Following antibody staining, cells were washed and propidium iodide (PI; Invitrogen) was added to exclude dead cells. Fluorescence activated cell sorting (FACS) was performed on the FACS Aria II (BD Biosciences) to isolate CD31^-^, CD45^-^, Sca1^-^, α7-integrin^+^, and Vcam1^+^ MuSCs.

### Culture of freshly isolated MuSCs on defined stiffness substrates

Freshly isolated MuSCs were immediately plated onto ECM (Sigma-Aldrich) coated 96-well plates with defined substrate elasticities- 0.5kPa, 16kPa and 32kPa (CytoSoft). Cells were cultured for 10 hours at 37°C in a humidified incubator with 5% CO₂, followed by fixation and immunofluorescence staining for the indicated protein markers.

### Isolation and culture of single myofibers

Single myofibers were isolated from the extensor digitorum longus (EDL) muscle. Briefly, EDL muscles were dissected from the hindlimb of mice and digested in DMEM containing 0.2% Collagenase Type 1 (Worthington) for 90 minutes in a shaking water bath at 37°C. Following digestion, EDL muscles were gently triturated using a horse serum coated glass pipette to isolate intact single myofibers containing resident MuSCs. The single myofibers were fixed immediately in 4% paraformaldehyde (PFA) for 10 minutes (t0; time point 0).

### Pharmacological Inhibition of single myofibers

Single myofibers isolated from WT mice were cultured in DMEM supplemented with 10% horse serum in the presence of either vehicle control (PBS; 0 µg/mL) or the RhoA inhibitor, Rho1 (0.5 or 1 µg/mL; Cytoskeleton Inc.) for 8 hours (t8).

To inhibit SP1 activity, single myofibers isolated from WT mice were treated with the SP1 inhibitor Mitramycin A (Cayman Chemical). Mitramycin A was initially dissolved in DMSO to generate a 1 mM stock solution and subsequently diluted in PBS to a working concentration of 100 µM. Myofibers were treated with vehicle control (90% DMSO and 10% PBS; 0 nM) or Mitramycin A (30 nM or 100 nM) for 8 hours (t8). Following treatment, myofibers were fixed in 4% PFA for 10 minutes prior to analysis.

### Immunofluorescence staining

Fixed single myofibers or freshly isolated FACS purified MuSCs were permeabilized in PBS containing 0.2% Triton X-100 (PBST) for 10 minutes at room temperature. Samples were then blocked in PBST containing 10% goat serum for 30 minutes, followed by incubation with primary antibodies overnight at 4°C. Primary antibodies used in this study were: Pax7 (DSHB), phospho-MLC (Thr18, Ser19; Invitrogen), phospho-S6 (Ser235, 236; Cell Signaling), MyoD (Invitrogen), YAP1 (Cell Signaling), Dnmt1 (Cell Signaling), Dnmt3A (Epigentek), Dnmt3B (Epigentek), Tet1 (Epigentek), Tet2 (Epigentek), Tet3 (Epigentek), SP1 (Abcam), phospho-SP1 (Thr453; Invitrogen), and Cleaved Caspase 3 (Cell Signaling). The following day, samples were washed with PBST and incubated with fluorophore-conjugated secondary antibodies together with DAPI (Invitrogen) for nuclear staining. Secondary antibodies used in this study were: Alexa Fluor 546 goat anti-mouse, Alexa Fluor 488 goat anti-rabbit antibodies, Alexa Fluor 488 goat anti-mouse, and Alexa Fluor 546 goat anti-rabbit antibodies (Invitrogen). F-actin was visualized using Alexa Fluor 488-conjugated phalloidin (Invitrogen). After final washes, samples were mounted, and images acquired using the Leica Thunder Imager, Olympus BX63 fluorescence microscope, and Olympus (Evident) FV4000 confocal microscope. The mean protein expression in MuSCs were quantified using Fiji software. Graphs were generated using GraphPad Prism version 11.0.0.

### Statistical Analysis

Statistical analyses were performed using GraphPad Prism. Data are presented as mean ± SD unless otherwise stated. Comparisons between two groups were performed using paired two-tailed Student’s *t*-tests. Differences were considered statistically significant at *p* < 0.05. Biological replicate numbers are provided in the figure legends.

### Analysis of MuSC Area and Circularity

Single myofibers stained for phospho-S6 were used to quantify MuSC area and circularity. Image analysis was performed using Fiji. Regions of interest (ROIs) were manually drawn around the cytoplasmic pS6 signal to define individual MuSC boundaries. Cell area and circularity index were subsequently measured for each cell. Nuclear area was determined by drawing ROIs around DAPI stained nuclei of MuSCs.

### Active RhoA assay

FACS-isolated MuSCs were fixed in 4% paraformaldehyde (PFA) for 25 minutes, washed with PBS, and permeabilized in 0.05% Triton X-100 in PBS for 10 minutes at room temperature. Cells were then blocked in 5% FBS/PBS and incubated for 1 hour at room temperature with GST-tagged Rhotekin Rho-binding domain (RBD) protein (Cytoskeleton Inc.), which specifically binds active RhoA. Following PBS washes, cells were incubated with fluorophore-conjugated anti-GST antibody (Invitrogen) together with DAPI (Invitrogen) for 1 hour at room temperature. Cells were subsequently washed with PBS, mounted, imaged, and analyzed for mean intensity.

### Reverse Transcription and Quantitative Polymerase Chain Reaction (RT-qPCR)

Freshly isolated MuSCs were collected directly in TRIzol^TM^ reagent (Invitrogen), and total RNA was extracted using phenol-chloroform separation followed by isopropanol precipitation in the presence of GlycoBlue^TM^ coprecipitant (Invitrogen). Isolated RNA was treated with the TURBO DNA-free^TM^ (Invitrogen) reagent to remove genomic DNA contamination. cDNA was synthesized from purified RNA using the SuperScript^TM^ First-Strand Synthesis System (Invitrogen) according to the manufacturer’s instructions. For quantitative RT-qPCR analysis, 5ng of cDNA was used per reaction to assess gene expression levels using the QuantStudio 3 Real-Time PCR system (Thermo Fisher Scientific). Expression levels were normalized to GAPDH, and fold change relative to control samples was determined using the ΔΔCt method.

### RNA sequencing (RNA-seq)

RNA quality control, library preparation, sequencing, and primary data processing were performed by the Yale Center for Genome Analysis (YCGA). Total RNA quality and purity were assessed by measuring A260/A280 and A260/A230 ratios by Nanodrop. RNA integrity was evaluated using the Agilent Bioanalyzer. Samples with an RNA integrity number (RIN) ≥ 7 were used for library preparation. RNA-seq libraries were generated using the NEBNext Single Cell/Low Input RNA Library Prep Kit for Illumina (New England Biolabs) with 200 pg-50 ng of input RNA. cDNA synthesis was performed using a template-switching approach, followed by fragmentation, end repair, dA-tailing, adapter ligation, and PCR amplification using the NEB Ultra II FS workflow. Indexed libraries passing quality and quantity thresholds were quantified by qPCR using the KAPA Library Quantification Kit and analyzed for insert size distribution using either the LabChip GX or Agilent Bioanalyzer platform. Libraries with yields ≥ 0.5 ng/µL were advanced for sequencing. Libraries were normalized to 2.0 nM and sequenced on the Illumina NovaSeq platform using 100 bp paired-end reads according to Illumina protocols. A 0.3% PhiX spike-in control was included in each sequencing lane to monitor sequencing quality. Sequencing data were processed using Illumina Real Time Analysis (RTA) software for base calling and demultiplexing. Primary analysis was performed using the Illumina CASAVA 1.8.2 software suite.

### RNA-seq data analysis

Raw and trimmed read quality was assessed using FastQC v0.12.11.^87^ Adapter trimming and removal of overrepresented sequences were performed using Cutadapt v4.4.^88^ Reads were aligned to the mouse reference genome (GRCm39/mm39) using STAR v2.7.11b^89^ and subsequently sorted and indexed using SAMtools v1.21.^90^ Gene-level read quantification was performed using featureCounts v1.6.4.^91^ Count matrices were imported into RStudio v2023.6.0.421, and differential gene expression analysis was carried out using DESeq2 v1.34.0.^92^ Differentially expressed genes were identified using a significance threshold of p ≤ 0.05 and an absolute fold change > 1.5. Gene ontology (GO) enrichment analysis of biological processes for both upregulated and downregulated genes was performed using clusterProfiler v4.2.2,^93^ and selected significant GO terms were plotted.

### Alternative Splicing Analysis

Aligned RNA-seq reads generated for downstream expression analysis were used for alternative splicing analysis. Reads were aligned to the mouse reference genome (GRCm39/mm39). Differential alternative splicing events were identified using rMATS v4.3.0.^94^ Five classes of alternative splicing events were analyzed, including skipped exons (SE), mutually exclusive exons (MXE), alternative 5ʹ splice sites (A5SS), alternative 3ʹ splice sites (A3SS), and retained introns (RI). rMATS output files (MATS.JC.txt) were imported into RStudio v2023.6.0.421 for downstream processing and visualization. Differential splicing events were considered significant at a false discovery rate (FDR) < 0.05 and an absolute percent spliced-in difference (|ΔPSI|) > 0.1.

### Chromatin Immunoprecipitation (ChIP) and qPCR

ChIP experiments were performed using 1×10^5^ FACS isolated MuSCs following the Zymo-Spin^TM^ ChIP kit (Zymo Research) protocol according to the manufacturer’s instructions. Briefly, cells were crosslinked with formaldehyde at a final concentration of 1% (v/v), and the crosslinking reaction was quenched by adding glycine to a final concentration of 0.125 M. Cells were washed with chilled 1× PBS containing 1 mM phenylmethylsulfonyl fluoride (PMSF) and 1× protease inhibitor cocktail (PIC). Crosslinked cell pellets were resuspended in chilled Nuclei Prep Buffer containing 1 mM PMSF and 1× PIC and incubated on ice for 5 minutes. Following centrifugation at 3,000 × g for 1 minute at 4°C, nuclei were resuspended in chilled Chromatin Shearing Buffer supplemented with 1 mM PMSF and 1× PIC, and incubated on ice for an additional 5 minutes. Chromatin was then sonicated for 5 minutes (Peak power-75, Duty factor-10, Cycles- burst 200) using Covaris M220 focused ultrasonicator. Sheared chromatin was incubated overnight at 4°C with either rabbit IgG (Invitrogen) or with SP1 antibody (Abcam) under rotation. Immunocomplexes were captured using ZymoMag Protein A beads for 1 hour at 4°C with rotation. Bead-bound chromatin complexes were sequentially washed with Chromatin Wash Buffers I, II, and III, with magnetic separation between washes. For elution and reverse crosslinking, bead-bound chromatin was resuspended in Chromatin Elution Buffer containing NaCl and incubated at 75°C followed by reverse crosslinking at 65°C. Samples were subsequently treated with Proteinase K. Immunoprecipitated DNA fragments were purified by phenol-chloroform extraction. Quantitative PCR analysis was performed using SYBR Green (Invitrogen) on QuantStudio 3 Real-Time PCR system (Thermo Fisher Scientific). SP1 ChIP signal was normalized to IgG control, and fold enrichment was determined using the ΔΔCt method.

### Isolation of Genomic DNA

Genomic DNA was isolated using a phenol-chloroform extraction method. Samples were centrifuged at 2,000 rpm for 5 minutes to pellet the cells. Sterile 1× PBS was added to the cell pellet to resuspend the cells and an equal volume of phenol/chloroform/isoamyl alcohol is added. Samples were mixed vigorously by inversion and centrifuged at 17,000 × g for 5 minutes at room temperature. The aqueous phase was transferred to a fresh tube, and DNA was precipitated by the addition of GlycoBlue^TM^ (Invitrogen), 7.5 M ammonium acetate, and 100% ethanol, followed by overnight incubation at - 20°C. The following day, samples were centrifuged at maximum speed for 30 minutes at 4°C to pellet the DNA. Pellets were washed with 75% ethanol, centrifuged again for 5 minutes at 4°C, air-dried for 5-10 minutes at room temperature, and resuspended in nuclease-free water. DNA concentration was quantified, and purified DNA was stored at -20°C until further use.

### DNA Methylation (Enzymatic Methylation; EM) Sequencing

Genomic DNA quality control, library preparation, sequencing, and primary data processing were performed by the Yale Center for Genome Analysis (YCGA). Genomic DNA samples underwent stringent quality control assessment, including quantification, purity analysis by A260/280 and A260/230 absorbance ratios, and 1% agarose gel electrophoresis to confirm DNA integrity and absence of RNA contamination. Indexed paired-end whole-genome methylation sequencing libraries were generated using the NEBNext Enzymatic Methyl-seq Library Kit (New England Biolabs). Briefly, 10–20 ng of genomic DNA was sheared to an average fragment size of 240-290 bp using a Covaris E220 focused ultrasonicator. Fragment size distribution was assessed using the TapeStation 4200 (Agilent). Fragmented DNA underwent end repair, adapter ligation, enzymatic oxidation of 5-methylcytosine and 5-hydroxymethylcytosine, cytosine deamination, and PCR amplification according to the manufacturer’s instructions. Dual-indexed libraries were quantified by qPCR using the KAPA Library Quantification Kit and assessed for insert size distribution on the TapeStation 4200. Libraries with yields ≥2 ng/µL were advanced for sequencing. Libraries were normalized to 2 nM and sequenced on the Illumina NovaSeq 6000 platform using 151 bp paired-end reads according to Illumina protocols. A 1% PhiX spike-in control was included in each sequencing lane for real-time quality monitoring. Sequencing data were processed using Illumina Real Time Analysis (RTA) software for base calling and demultiplexing. Primary sequence analysis and alignment were performed using the Illumina CASAVA 1.8.2 software suite.

### EM-seq data analysis

Differentially methylated regions (DMRs) were identified from EM-seq data using a multi-step analysis pipeline. Sequencing reads were aligned to the mouse reference genome (GRCm39) using Bismark (v0.24.2), which supports mapping and methylation calling for EM-seq libraries in a manner analogous to bisulfite sequencing–based workflows. Cytosine methylation calls were then generated with bismark_methylation_extractor, producing genome-wide cytosine methylation reports for each sample. These reports were processed with DMRfinder (v0.3),^95^ first using the combine_CpG_sites.py script to cluster nearby CpG sites into candidate regions with the following parameters: -r 10, -s 4, -d 100, -c 3, -x 500, and -m 20. Finally, the findDMRs.r script was applied to these regions to perform statistical testing and identify significant differentially methylated regions between experimental groups.

## RESOURCE AVAILABILITY

### Lead contact

Requests for further information and resources should be directed to and will be fulfilled by the lead contact, Susan Eliazer.

### Materials Availability

This study did not generate new unique reagents.

### Data and code availability

- RNA sequencing (GSE329437) and Enzymatic Methylation sequencing (GSE329791) datasets from skeletal muscle stem cells have been deposited at Gene Express Omnibus (GEO) and are publicly available on May 31^st^, 2026.
- This paper does not report original code.
- Any additional information required to reanalyze the data reported in this paper is available from the lead contact upon request

## ACKNOWLEDGMENTS

We would like to thank Drs. Charles Keller, Cord Brakebusch, and Maria Behrens for providing the mice used in this study. We also thank the University of North Dakota (UND) Epigenetics Group faculty members, Drs. Archana Dhasarathy, Sergei Nechev, and Motoki Takaku, for their valuable feedback and suggestions on the manuscript. We are grateful to the members of the Eliazer laboratory for their insightful discussions and contributions during the preparation of this manuscript. We acknowledge the technical assistance provided by the UND Flow Cytometry (FACS) Core, supported by NIH grant P20GM113123; the UND School of Medicine and Health Sciences (SMHS) Imaging Core, supported by NIH grant P20GM113123, DaCCoTA CTR NIH grant U54GM128729, and UND SMHS funds; and the UND Pathology Imaging Core, supported by an Institutional Development Award (IDeA) from the NIH/NIGMS under grant number P20GM103442. This work was supported by the NIH/NIGMS Epigenetics CoBRE grant P20GM104360 to S.E. and by the University of North Dakota Vice President for Research (VPR) Postdoctoral Seed Funding to P.J.

## AUTHOR CONTRIBUTIONS

P.J. designed and performed experiments, analyzed data, interpreted results, and wrote the manuscript. P.R.D. performed experiments, and analyzed data. J.P. performed experiments, and analyzed data. C.O.A. analyzed data, and interpreted results. A.G. performed experiments. S.A. performed experiments. C.W.D.J. performed experiments. D.P. analyzed data, and interpreted results. S.E. conceptualized the project, designed and interpreted experiments, and wrote the manuscript.

